# Clustering based approach for population level identification of condition-associated T-cell receptor β-chain CDR3 sequences

**DOI:** 10.1101/490102

**Authors:** Dawit A. Yohannes, Katri Kaukinen, Kalle Kurppa, Päivi Saavalainen, Dario Greco

**Author notes:** These authors contributed equally to this work (Päivi Saavalainen, Dario Greco).

## Abstract

**Motivation:** Deep immune receptor sequencing, Repseq, provides unprecedented opportunities to identify condition-associated T-cell clones, represented by T-cell receptor (TCR) CDR3 sequences. TCR profiling has potential value for increasing immunopathological understanding of various diseases, and holds considerable clinical relevance. However, due to the immense diversity of the immune repertoire, identification of condition relevant TCR CDR3s from total repertoires has so far been limited either to mostly “public” CDR3 sequences, which are shared across unrelated individuals, or to comparisons of CDR3 frequencies from multiple samples from the same individual. A methodology for the identification of condition-associated TCR CDR3s by population level comparison of groups of Repseq samples is currently lacking.

**Results:** We implemented a computational pipeline that allows population level comparison of Repseq sample groups at the level of the immune repertoire sub-units that are shared across individuals. These sub-units (or sub-repertoires) represent shared immuno-genomic features across individuals that potentially encode common signatures in the immune response to antigens. The method first performs unsupervised clustering of CDR3 sequences within each sample based on their similarity in nucleotide or amino acid subsequence frequency. Next, it finds matching clusters across samples, the immune sub-repertoires, and performs statistical differential abundance testing at the level of the identified sub-repertoires. We applied the method on total TCR CDR3β Repseq datasets of celiac disease patients in gluten exposed and unexposed conditions, as well as on public dataset of yellow fever vaccination volunteers before and after immunization. The method successfully identified condition-associated CDR3β sequences, as evidenced by considerable agreement of TRBV-gene and positional amino acid usage patterns in the detected CDR3β sequences with previously known CDR3β species relevant to celiac disease. The method also recovered significantly high numbers of previously known CDR3β sequences, relevant to each condition than would be expected by chance. We conclude that immune sub-repertoires of similar immuno-genomic features, shared across unrelated individuals, encode common immunological information. Moreover, they can serve as viable units of population level immune repertoire comparison, serving as proxy for identification of condition-associated CDR3 sequences.

## Introduction

Targeted high-throughput sequencing of T-cell receptors, Repseq, has enabled in-depth profiling of immune repertoires (Benichou *et al*., 2012). One critical application of Repseq technology is the identification of condition-associated T-cell clones based on observed changes in T-cell clone frequencies. This allows the tracking of immune cells that have expanded or contracted following antigen exposure or treatment. Such analysis, however, is complicated by the fact that T-cell receptor (TCR) sequences are highly diverse, with estimated tens of millions unique TCR expressing T-cell clones largely unique to individuals (Vanhanen *et al.*, 2016; Qi *et al.*, 2014), making direct comparison of T-cell clone abundances across multiple sample groups challenging.

A frequently used approach to the identification of condition-specific clonotypes across sample groups is the investigation of the so called public clonotypes (represented typically by unique TCR CDR3 sequences), which are commonly observed across many individuals (Venturi *et al.*, 2008; Li *et al.*, 2012; Benati *et al.*, 2016; Covacu *et al.*, 2016; Madi *et al.*, 2014; Emerson *et al.*, 2017; Pogorelyy, Minervina, Chudakov, *et al.*, 2018). However, such shared clonotypes make up a small portion of the total immune response in each individual, and more should be learned about the adaptive immune response by also studying the private response. For instance, others and we have found that only around 10% of the response to gluten in celiac disease (CD) patients involves public CDR3 sequences (Yohannes *et al.*, 2017; Risnes *et al.*, 2018). On the other hand, DeWitt *et al.* compared frequencies of clonotypes in repertoires sampled from the same individual to identify differentially abundant clonotypes, thus identifying disease-relevant clonotypes within each individual (DeWitt *et al.*, 2015). This approach allows the identification of interesting clones, which are mostly private to individuals, but does not allow the investigation of differentially abundant clonotypes at the population level. Moreover, it requires acquisition of multiple samples from each individual. Thus, there is currently a need for population level differential abundance analysis methods for the identification of condition-specific T-cell clonotypes from Repseq data.

We recently showed that over-represented amino acid motifs in CD-associated TCR CDR3β sequences, originally identified from tetramer binding antigen-reactive T-cells (Qiao *et al.*, 2011, 2013), were also detectable from the unsorted total peripheral blood immune repertoires of celiac disease patients despite the immense repertoire diversity. This observation and closer inspection of the CD-associated TCR CDR3β sequences, strongly suggested that CDR3β sequences associated to CD exhibit sequence-level similarities that can be used to group them into clusters, reflecting similar immunogenomic features involved in the immune reaction that are possibly shared by patients. Such high sequence similarity had been observed in B-cell receptors (BCRs) associated to chronic lymphocytic leukemia (referred to as BCR stereotypy), with particular relevance in patient stratification of clinical relevance (Agathangelidis *et al.*, 2012; Darzentas and Stamatopoulos, 2013). More recently, Dash *et al.* and Glanville *et al.* have shown that antigen-specific TCR sequences collected from different patients could be clustered into antigen specificity groups that share sequence similarity (Dash *et al.*, 2017; Glanville *et al.*, 2017). Overall, recent Repseq studies have reported sequence similarity in CDR3 sequences associated with conditions, a characteristic of the immune response that could be harnessed for the identification of condition relevant CDR3 sequences from total unselected repertoires by comparing sample groups.

In this work, we propose differential abundance analysis at the level of shared clusters of T-cell receptor CDR3β sequences, called sub-repertoires, to enable identification of disease relevant clusters of CDR3β sequences by comparing Repseq experimental groups. We first applied within-sample CDR3β clustering to reduce the diversity of immune repertoires into manageable and comparable units of analysis. We showed that these clusters of receptors, made up of clonotypes with highly similar frequencies of nucleotide or amino acid subsequences (k-mers), form biologically meaningful units of analysis since they are commonly present in repertoires of unrelated individuals. We then performed statistical differential abundance analysis at the level of these sub-repertoires for the identification of condition-specific CDR3β clonotypes. We also showed that this methodology allows successful detection of condition-associated CDR3 sequences, both private and public, from immune repertoire datasets of celiac disease patients, and yellow fever virus vaccination volunteers, by comparing groups of samples at the population level.

## Methods

Our methodology narrows down the highly diverse immune repertoire data and helps to identify disease associated differentially abundant CDR3 sequences by comparing samples from two treatment groups. As input, it requires high-throughput T-cell receptor CDR3 sequences (of either CDR3α or CDR3β), pre-processed for each sample into a format that is usually tab-delimited text file, with each CDR3 and its features (its Nucleotide & Amino acid sequences, CDR3 length, V-,D-,and J-gene segment usage, etc.) in one row. Referring to each repertoire data simply as a sample, the method processes the samples in four major steps to identify differentially abundant CDR3 sequences (see Figure 1):

**Figure 1:**
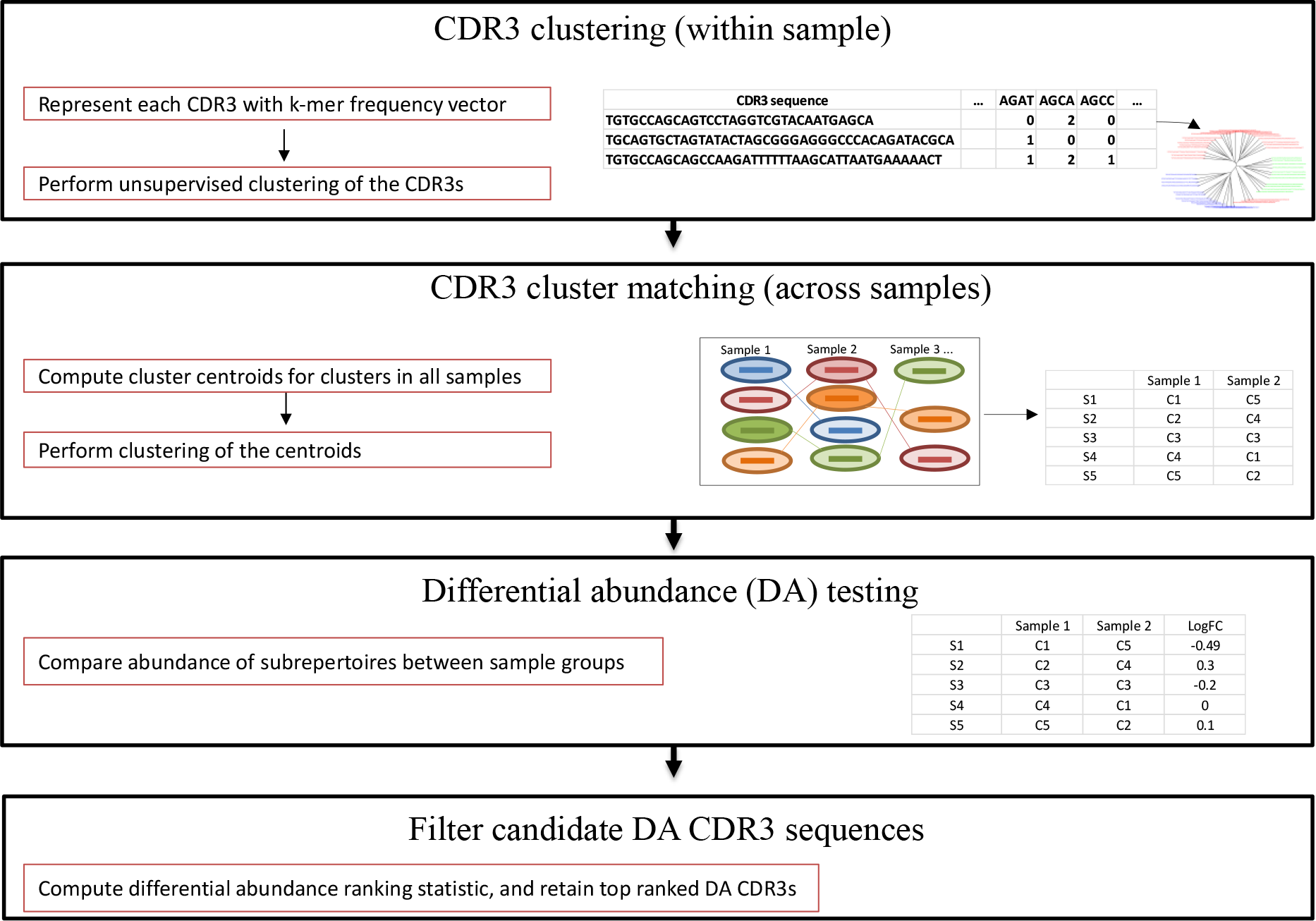
Schematic of the clustering-based differential abundance detection methodology using CDR3 repertoire HTS data. In the first step, CDR3 sequences are first clustered within sample (agglomerative hierarchical clustering with complete linkage method), using a pairwise distance matrix computed from nucleotide or amino acid subsequence or k-mer frequency vectors for each CDR3 (typically k-mer=4 for nucleotide, k-mer=3 for amino acid level analysis). Next, cluster of CDR3s are defined. In the second step, we look for matching clusters across samples by clustering their centroids. The cluster centroids in all samples are collected and fed to K-means (with k determined as discussed in method section), a cluster match table is produced which lists the label of the matching clusters in each repertoire sample, representing shared immunogenomic information in all the samples. In the third step, differential abundance testing is performed at the sub-repertoire level between groups of samples. If differentially abundant (DA) sub-repertoires are detected, filtering of the CDR3s belonging to the DA sub-repertoires is performed to identify CDR3s with statistically significant association to the condition under study.

1. CDR3 clustering: each CDR3 in a sample is first represented using a high dimensional k-mer frequency vector by counting the frequency of each possible contiguous nucleotide (nt) or amino acid (aa) subsequences in the CDR3. We have typically used k=4 for nt k-mers, resulting in a feature vector size of 4^k = 256, or k=3 for aa k-mers resulting in a feature vector size of 20^3 = 8000. Next, the high-dimensional k-mer frequency vectors are used to perform unsupervised clustering of the CDR3 sequences within each sample (using agglomerative hierarchical clustering, with the complete linkage method which showed better cluster stability with more number of clusters-or “zooming-in”, Supplementary Figure 4S). We use the euclidean distance measure when using nt k-mer and the cosine distance when using aa k-mer vectors to determine the distance between a pair of CDR3s. After the hierarchical clustering, the dynamic tree cut algorithm is used to define the CDR3 clusters in each sample (Langfelder *et al.*, 2008).
2. Cluster matching across samples: For each cluster in each sample, the average frequency for each k-mer is computed from the members of the cluster to get the cluster centroid. The centroids of clusters from all samples are collected together, and unsupervised clustering of the centroids is performed using either hierarchical clustering or K-means to group centroids based on closeness in k-mer frequency profiles. When using k-means, clustering of the centroids is performed using k that has the maximum optimal-k (oK) score:

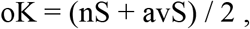

where nS is the proportion of clusters with silhouette value greater than the average silhouette (over all clusters), and avS is the average silhouette shifted to be between 0 and 1, by adding 1 and dividing by 2. oK values range from 0 to 1. oK is computed from all k starting from the minimum to the maximum number of CDR3 clusters per sample observed across all samples (from the result of step 1) to determine the k with maximum oK score. Next, each cluster of centroids is examined and, if multiple centroids from one sample are clustered together, the clusters of such centroids are merged in the original sample and the centroid updated. Otherwise, all clusters of centroids representing matching clusters from multiple samples (not necessarily all samples) are retained. Given N samples, this step generates a cluster match table with oK rows and N columns, in which row entries represent CDR3 cluster labels from all samples that are close (or have matching) centroids, representing underlying features encoding conserved immunological features in all N or some of the samples (matching clusters may not be found in all samples). We refer to each row in this cluster match table as a sub-repertoire.
3. Differential abundance testing: sub-repertoires that exist in at least x number of samples per group are first selected. A sub-repertoire abundance matrix is then generated for the selected sub-repertoires, which contains the abundance of each sub-repertoire in each sample. This sub-repertoire abundance matrix can be generated in different ways. Typically, the original samples are first normalized to same total CDR3 sizes (counts). Then, the abundance of a sub-repertoire in a sample is calculated as the sum of CDR3 counts belonging to that sub-repertoire in the sample; or the relative frequency of that sum is used as a relative abundance estimate; or to avoid bias in the previous two abundance estimates that might arise due to differences in the number of CDR3s per sub-repertoire in a sample, the relative clone size in sub-repertoires is used as a proxy for abundance (i.e., average clone count in sub-repertoire / average clone count in sample). Next, for each sub-repertoire, differential abundance testing between the two groups of samples is performed using various tests. We used paired t-test, and two-class, unpaired, RankProd (Breitling *et al.*, 2004) test, both of which work well. The CDR3 sequences belonging to significantly differentially abundant (DA) sub-repertoires are then extracted from each sample; these are candidate DA CDR3 sequences.
4. Filtering candidate DA CDR3s: DA sub-repertoires are not “pure” and contain CDR3s that are not necessarily associated with the condition thus requiring further filtering. To do this, all candidate DA CDR3 sequences (i.e., all CDR3 sequences that are in DA sub-repertoires) are first ranked as follows. For each candidate CDR3 *i*, a rank sum, C*i*, is computed by adding the candidate’s ranks from 6 factors:

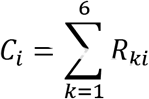

where R_1*i*_ is candidate *i*’s importance in classifying the groups (random forest mean decrease in accuracy (Ho, 1995; Liaw and Wiener, 2002)), R_2*i*_ is its mean fisher’s exact test p-value calculated by comparing its abundance in each paired samples separately, R_3*i*_ is its mean odds ratio from fisher’s exact test calculated by comparing its abundance in each paired samples separately, R_4*i*_ is its mean nucleotide to amino acid (nt-to-aa) ratio across the samples to account for its level of convergent selection, R_5*i*_ is the difference in the number of samples per condition in which it exists, to account for its degree of condition induced “public-ness”, and R_6*i*_ is the number of times it has been detected in repeat resample runs of the DA analysis (Repseq datasets are huge datasets and require a lot of computational resources, we thus perform repeat sub-sampling of the raw datasets and run the differential abundance analysis of steps 1-3 for each resampled datasets, the candidate DA CDR3 sequences from each round are collected, CDR3 sequences that have been detected as candidate in multiple repeat resamples are given higher rank, i.e., rank of 1). The ranking in each factor is defined differently for assessing enrichment and de-enrichment. When assessing enrichment highest rank is given for highest mean decrease in accuracy for R_1_, smallest p-value for R_2_, highest odds ratio for R_3_, highest nt-to-aa ratio for R_4_, highest increase in detection in the condition group for R_5_, and highest number of detection in multiple runs of the analysis for R_6_. For assessment of de-enrichment, highest rank is given for lowest mean decrease in accuracy for R_1_, smallest p-value for R_2_, lowest odds ratio for R_3_, lowest nt-to-aa ratio for R_4_, highest decrease in detection in the condition group for R_5_, and lowest number of detection in multiple runs of the analysis for R_6_. The minimum value of 1 in each factor R signifies high rank. Since the range of rank values is different for each rank type, all Rs are scaled to be between 0 and 1 by subtracting the minimum and dividing by the range. We then calculate the p-value for C_*i*_ using a randomization test, as the proportion of n rank sum values, calculated from n permutations (random shuffling of all Rs from the 6 factors, we typically used n = 1000), that are equal or less than C_*i*_. We consider candidate DA CDR3s with C p-values less than 0.05 and q-value (minimal FDR at each p-value) less than 0.05 as differentially abundant CDR3s. All six ranking factors were given equal weight in our analysis in the calculation of the rank sum C, but different weights could be used for each factor depending on the application. For false discovery rate estimation, we used a decoy-based strategy by including in the analysis randomly drawn CDR3 sequences from a reference database of healthy TCR CDR3 PBMC repertoires to each sample’s repertoire data. The rate of false detection of the decoy CDR3s was used to estimate the FDR and q-value at each p-value level for all candidate CDR3s ordered from smallest to highest C p-values.

We applied the method in the following manner for the TCR CDR3β repertoire datasets in this work: 1) 10 runs of steps 1 to 3 of the pipeline using randomly selected subsamples of 5000 unique CDR3β sequences for each sample. 2) Within sample CDR3β clustering using either nucleotide 4-mers or amino acid 3-mer feature vectors. 3) sub-repertoire level differential abundance detection using paired t-test with the p-value cut-off of 0.1, as we found from evaluations of results from multiple runs that the less stringent p-value cut-off for comparing sub-repertoire abundances across samples helps increase the signal to noise ratio by providing the right level of “zooming-in”, without compromising detection capacity in downstream steps of the pipeline. 4) Combining the candidate CDR3βs from the 10 runs and performing the filtering step, CDR3βs with p-value and q-value less than 0.05 were then considered condition-associated CDR3β sequences.

### Characterization of differentially abundant CDR3β sequences

To evaluate TRBV gene usage and per position amino acid usage among differentially enriched CDR3β sequences, we compared the observed usage frequencies in the list of enriched CDR3βs to the frequencies obtained from 100 randomly sampled sets of CDR3βs (same size as the enriched list) from the combined dataset off all samples. The significance of the observed frequencies was calculated as the proportion of frequencies in the randomly sampled sets that were equal or more than the observed frequencies (p-values less than 0.05 were considered statistically significant).

### *Enrichment analysis of known condition-associated* CDR3β_*s*_

To evaluate the effectiveness of the method in identifying CDR3βs that are truly associated with a condition, we compared the number of observed CDR3βs known to be associated with the condition (from previous studies), to the frequencies of such known CDR3βs obtained from 100 random sets of CDR3βs (same size as the enriched list) sampled from the combined dataset of all samples. The significance of the observed frequency of known CDR3βs was calculated as the proportion of known CDR3βs in the randomly sampled sets that were equal or more than the observed frequency. Alternatively, the proportion of known CDR3βs in the list of enriched CDR3βs was compared to the proportion of known CDR3βs in the total combined repertoire of samples in a dataset using fisher’s exact test with p-values less than 0.05 considered statistically significant.

### Datasets

We used three TCR CDR3β immune repertoire datasets to test the method (Table 1). Celiac disease (CD) PBMC and Gut datasets of our celiac disease study cohort (Yohannes *et al.*, 2017), and yellow fever vaccination (YFV) PBMC dataset (DeWitt *et al.*, 2015) obtained from the public immune repertoire database, immuneACCESS, of Adaptive Biotechnologies (immuneACCESS, Adaptive Biotechnologies, Seattle, WA. Available from: http://adaptivebiotech.com/pub/dewitt-2015-jvi, Accessed on June 11, 2018). The sample preparation and sequencing of the datasets have been described before (Yohannes *et al.*, 2017; DeWitt *et al.*, 2015); in brief, genomic DNA was extracted from total PBMC for Celiac disease patients in CD PBMC dataset, and yellow fever virus vaccination volunteers in YFV PBMC dataset, or from gut biopsy for celiac disease patients in CD Gut dataset. The TCR CDR3β region was then deeply sequenced using the ImmunoSeq assay (Robins *et al.*, 2009) (www.adaptivebiotech.com; www.immunoseq.com), which uses optimized multiplex PCR to amplify the TCR CDR3β region, and Illumina for sequencing. It then determines the CDR3β sequences and their abundance, as well as annotates the gene segments according to International ImMunoGeneTics (IMGT) specifications (Lefranc *et al.*, 2015).

**Table 1:** The three datasets used in the study. The number of total and unique nucleotide CDR3β sequences is shown for each sample.

| Dataset | Sample name | Group 1 |  | Group 2 |  |
| --- | --- | --- | --- | --- | --- |
| Celiac Disease (CD) PBMC (n=4) |  | Day 0 (before eating wheat gluten) |  | Day 6 (After eating wheat gluten for 3 days) |  |
|  |  | Productive Total CDR3 | Productive unique CDR3 | Productive Total CDR3 | Productive unique CDR3 |
|  | CD005 | 527382 | 16174 | 589603 | 24195 |
|  | CD006 | 794514 | 13013 | 582690 | 15179 |
|  | CD011 | 541814 | 16014 | 384155 | 10452 |
|  | CD039 | 339279 | 10221 | 436176 | 10403 |
| Celiac Disease (CD) Gut (n=5) |  | GFD (after 1 year gluten free diet treatment) |  | ACT-disease (Sampled right after diagnosis) |  |
|  |  | Productive Total CDR3 | Productive unique CDR3 | Productive Total CDR3 | Productive unique CDR3 |
|  | CD1GB | 1189542 | 5886 | 1233903 | 10434 |
|  | CD2GB | 1760963 | 10033 | 968548 | 11715 |
|  | CD3GB | 3339752 | 8284 | 1101307 | 9863 |
|  | CD4GB | 2197571 | 9567 | 757564 | 10423 |
|  | CD5GB | 761406 | 6266 | 830042 | 5370 |
| Yellow Fever Vaccine (YFV) immunization data (PBMC n=9) |  | Day 0 (pre-vaccination of single dose of YF-17D) |  | Day 14 (post-vaccination) |  |
|  |  | Productive Total | Productive unique | Productive Total | Productive unique |
|  | Subject 1 | 5520003 | 334966 | 9034146 | 365705 |
|  | Subject 2 | 7546126 | 385506 | 9535753 | 434828 |
|  | Subject 3 | 4351030 | 339330 | 8930117 | 348878 |
|  | Subject 4 | 4522888 | 244022 | 8343948 | 292385 |
|  | Subject 5 | 1281823 | 64751 | 9201942 | 316705 |
|  | Subject 6 | 4525600 | 350914 | 6873009 | 275193 |
|  | Subject 7 | 3672998 | 173178 | 25873400 | 222176 |
|  | Subject 8 | 4761179 | 277491 | 9991028 | 258036 |
|  | Subject 9 | 3533232 | 207966 | 9110762 | 228819 |

### Benchmarking

We compared the performance of the method to four recently published methods for celiac-associated CDR3 detection in the CD PBMC dataset. The methods are vdjRec (Pogorelyy, Minervina, Chudakov, *et al.*, 2018), DeWitt’s method (DeWitt *et al.*, 2015), and Alice (Pogorelyy, Minervina, Shugay, *et al.*, 2018), and our previously published method here referred to as the Yohannes method (Yohannes *et al.*, 2017). Since there is no exhaustive truth set of CD associated CDR3s, we compared the methods by how many of 56 previously published, well-known CD-associated CDR3s present in the total CD PBMC dataset that the methods detect. The 56 celiac disease associated CDR3s in our dataset are among those published by Qiao *et al.*, Han *et al.* and Petersen *et al.* as gluten tetramer reactive CDR3βs, thus are a suitable truth set for comparing the methods (Qiao *et al.*, 2011; Han *et al.*, 2013; Petersen *et al.*, 2014). Input data for vdjRec and Alice for every V-J combination in CD PBMC datasets was prepared, and estimation of CDR3 generation probability was performed as described by the methods. Both vdjRec and Alice were run only on the four gluten exposed day 6 samples of CD PBMC. DeWitt’s method was implemented in R and was run on each sample pair with minimum total count cutoff of 100 per CDR3.

### Data availability

The CD PBMC and CD Gut datasets are available from the corresponding author on reasonable request. The YFV PBMC dataset can be accessed from immuneACCESS (available at: http://adaptivebiotech.com/pub/dewitt-2015-jvi). R implementation of the method is available at https://github.com/Greco-Lab/RepAn.

## Results

We developed a computational pipeline that allows the detection of differentially abundant CDR3 sequences between groups of deeply sequenced immune repertoire datasets in two conditions. The method involves four major steps (Figure 1, see Method section for details). In the first two steps, CDR3 sequences in samples are divided into clusters of similar CDR3 sequences, which are then matched across samples to define sub-repertoires that exist in multiple samples possibly encoding similar immunological information. In the next two steps, statistical testing for differential abundance is performed at the defined sub-repertoire levels to identify condition-associated CDR3 sequences. Such an approach was initially inspired by previous reports that described linear amino acid motifs in the CDR3β sequences responsive to gluten in celiac disease (Qiao *et al.*, 2013; Han *et al.*, 2013). We also reported detecting such celiac disease associated public CDR3βs from the global immune repertoire without pre-selecting for antigen specific T-cell populations by simply comparing CDR3β abundances between different conditions (Yohannes *et al.*, 2017). We had generally observed that the CD associated CDR3β sequences could potentially be grouped into clusters based on sequence similarity, reflecting possible shared specificity in the adaptive immune response.

We hypothesized that grouping of CDR3s in the global repertoire could reduce the enormous diversity of the immune repertoires into manageable units, with the potential of allowing comparison of such CDR3 clusters between sample groups. To investigate the validity of such an approach, we first evaluated if a cluster of CDR3β sequences in one sample could be similar, in terms of subsequence composition, to another cluster in another sample. Importantly, the cluster must be closer to its match in another sample than it is to other clusters of CDR3βs in its home sample, ideally signifying conserved immunogenomic features in the samples.

Indeed, we observed that such clusters of CDR3βs, with closely similar subsequence composition, exist across samples even in unrelated individuals. For example, for the two CD PBMC repertoire samples CD005 and CD006 (unrelated celiac disease patients), we first subsampled 3000 CDR3βs each from their total unique nucleotide CDR3β sequences. Unsupervised clustering of the CDR3βs was then performed within each sample based on the pairwise distance of the CDR3βs in subsequence composition, and the clusters of CDR3βs in each sample was determined. The centroids of all clusters from both samples were then pooled, and clustered again to identify matching CDR3β cluster centroids (steps 1 and 2 on Figure 1). Out of the 27 identified centroid clusters i.e., sub-repertoires, 25 (∼93%) had centroids representing CDR3β clusters from both samples (Figure 2a), suggesting that the CDR3βs represented by such centroids encode possibly shared immunological information in the two samples. In other words, in such sub-repertoires, a cluster from one sample is closer to another CDR3β cluster in the second sample than it is to many other clusters in its home sample (Figure 2a). We wondered if the subsequence composition similarity in sub-repertoires containing centroids from both samples arises mainly from similar V-gene segment usage. Comparison of the TRBV-gene usage frequency distribution between matching clusters from the two samples, however, suggests V-gene usage similarity does not necessarily explain the subsequence composition similarity (Figure 2b), with almost 50% of the sub-repertoires containing matching clusters with significantly different V-gene usage frequency profiles.

**Figure 2:**
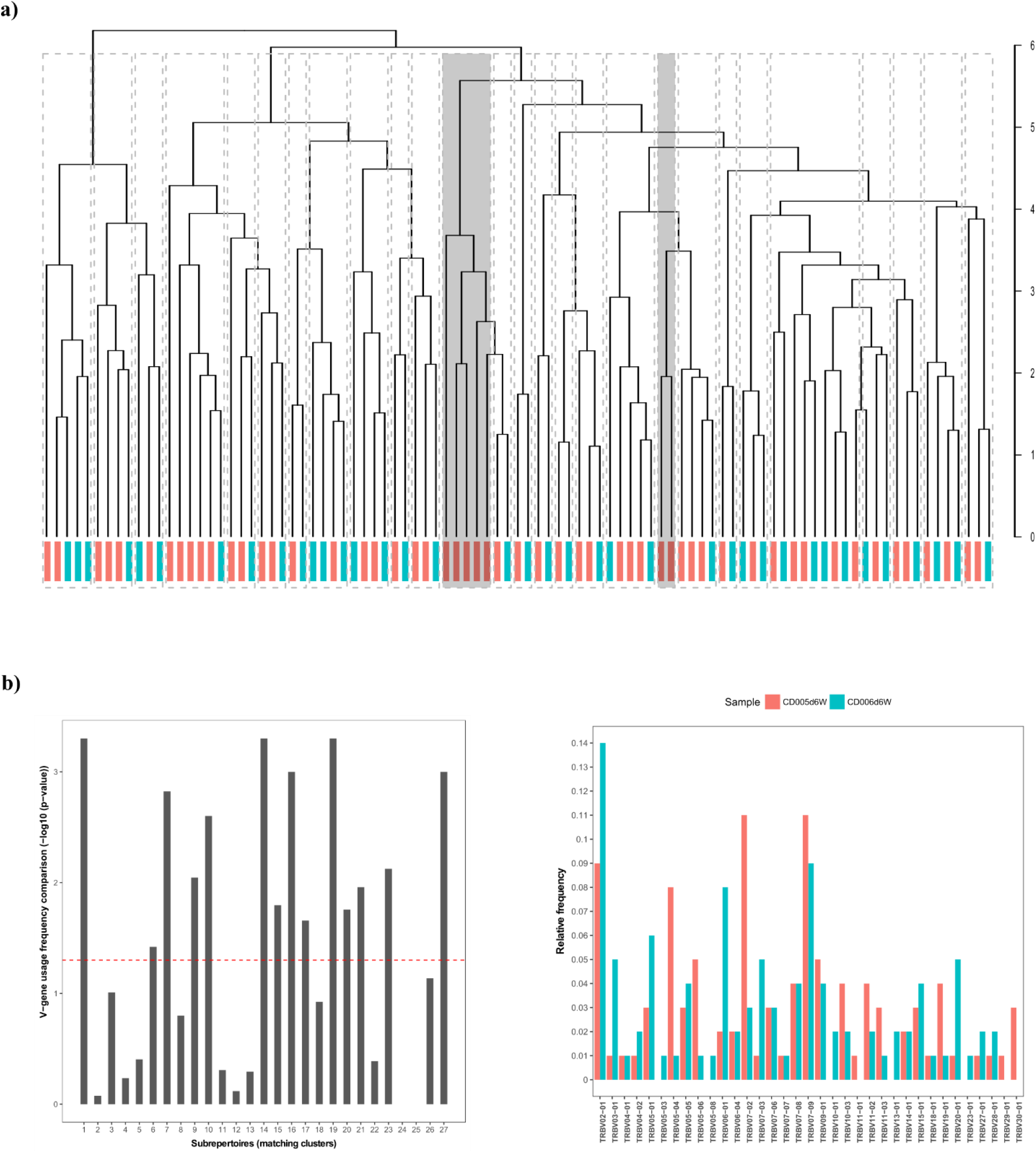
CDR3 sub-repertoire matching in samples of unrelated individuals. (a) hierarchical clustering of CDR3 cluster centroids from samples CD005 (red) and CD006 (green) from our CD PBMC dataset, identified 27 sub-repertoires of which 25 (93%) had cluster representatives from both samples. Only 2 of the 27 (7%) sub-repertoires (shown in grey) are homogenous, containing cluster centroids from only one sample. (b) For each sub-repertoire, v-gene usage frequency was compared between clusters coming from the two samples, the dotted line indicates the cut-off point at p-value 0.05 (using chi-square test of independence) in −log10(p-value) above which the v-gene usage is significantly different (left panel). An example v-gene usage profile is shown for clusters belonging to sub-repertoire 1 (right panel).

We then investigated the extent of sub-repertoire sharing when there are more number of samples in the analysis. We performed the clustering and cluster matching analysis on all 8 samples of our celiac disease (CD) PBMC datasets, from both pre-gluten challenge (day 0) and post-gluten challenge (day 6) conditions, at different sequencing depths per sample. Figure 3a shows proportions of sub-repertoires containing CDR3 cluster centroids from *n* number of the samples when the matching of centroids is performed using either hierarchical clustering (hc) or k-means (km), and the analysis performed at different repertoire sequencing depths. The proportion of sub-repertoires containing representative centroids from only one sample was negligibly low in all ten analyses using samples at different repertoire depths. An increasing trend was observed in the proportion of sub-repertoires containing representative centroids from *n* different samples as *n* increased, especially with k-means matching of cluster centroids, with the highest median in sub-repertoire proportions containing cluster centroids from across samples obtained when *n* is 8, i.e, from all samples. In the ten runs performed at different repertoire depths, the minimum proportions of sub-repertoires containing CDR3 cluster centroids from all 8 samples was approximately 10-20%. In addition, more than ∼40% (with hc matching) and 60% (with km matching) of the sub-repertoires contain centroids from at least 6 of the 8 samples (Figure 3b), suggesting that enough of the sub-repertoires are present in multiple samples to allow comparison of sub-repertoire abundance at the population level. Once the sub-repertoires that exist across samples are determined, statistical comparison of their abundance between sample groups is performed. If differentially abundant (DA) sub-repertoires are detected, further filtering of the CDR3s belonging to the DA sub-repertoires is performed to identify those with strong evidence of association to the condition under study (see Methods section for details).

**Figure 3:**
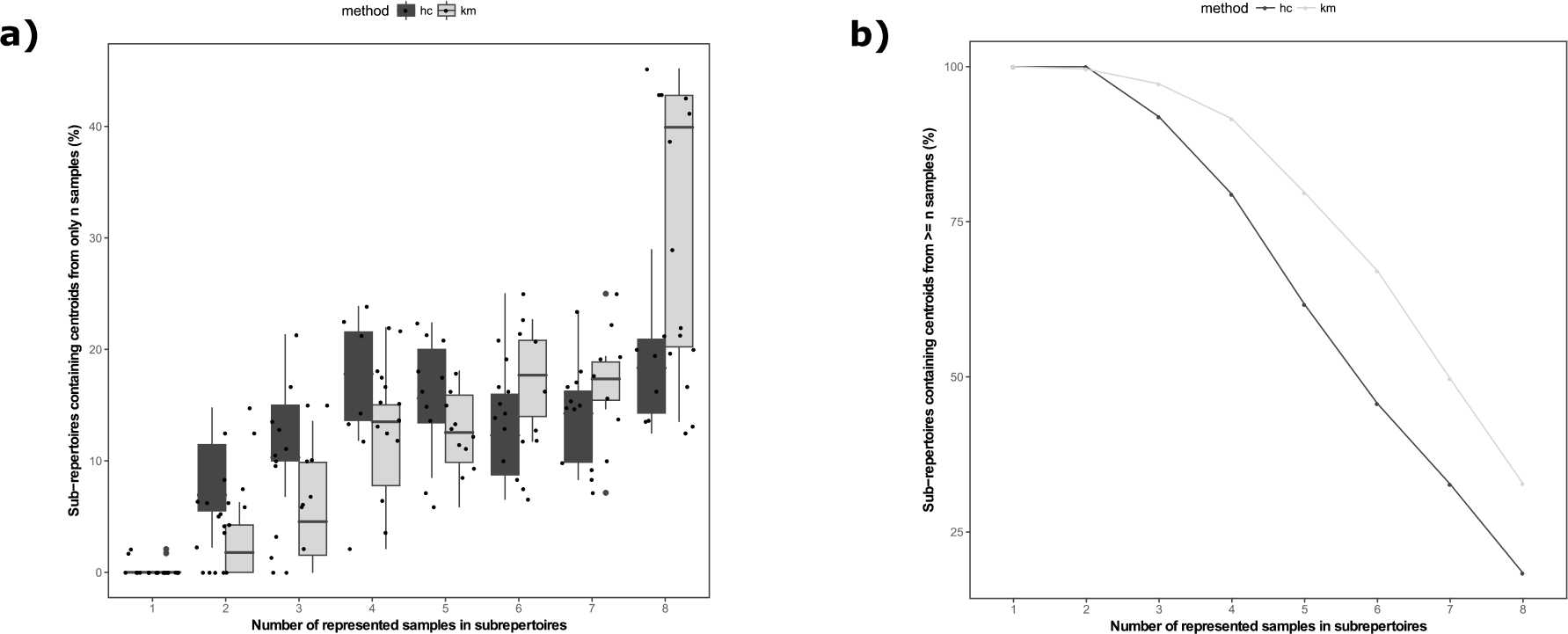
Sub-repertoire detection across many samples. CDR3 clustering, and sub-repertoire detection using both hierarchical clustering (hc) and k-means (km) clustering of the CDR3 cluster centroids was performed for all 8 CD PBMC samples. (a) Shows proportions of sub-repertoires containing CDR3 cluster centroids from only n samples, dots are the estimate from each of 10 analyses from subsampled repertoires with sequencing depths of 1 to 10 thousand unique nucleotide CDR3s per sample. (b) The cumulative proportion of sub-repertoires containing representative clusters from n samples or more is shown; the cumulative at each n is computed from the mean proportion of n represented samples from the 10 resample analyses.

We first applied the pipeline on our two high throughput T-cell CDR3β repertoire datasets of celiac disease patients, CD PBMC (n=4) and CD Gut (n=5), sampled from peripheral blood and gut biopsies respectively. We performed two analyses for each data, by clustering CDR3β sequences using nucleotide 4-mers and amino acid 3-mer feature vectors for assessing sequence composition at the nucleotide or amino acid levels respectively. For each analysis, we run the pipeline 10 times using randomly selected subsamples of 5000 unique CDR3β sequences for each sample and combined the results from all repeated resampling rounds for the CDR3β filtering step. Detecting differentially abundant (DA) sub-repertoires (step 3 in Figure 1) was done using paired t-test with the p-value cut-off of 0.1, and was applied by comparing sub-repertoire abundances only on sub-repertoires that exist in at least 3 samples per group. We found from evaluations of results from multiple runs, with combinations of the parameters, that the less stringent p-value cut-off for comparing sub-repertoire abundances across samples helps increase the signal to noise ratio for downstream steps in the pipeline, without compromising sensitivity, and giving us enough number of candidate CDR3β sequences for the filtering analysis in step 4. The filtering of the candidates was then performed and candidate CDR3βs with a p-value less than 0.05 were considered condition-associated CDR3β sequences.

For the CD PBMC dataset, the method identified 459 CDR3β sequences that showed significant enrichment and 441 that showed de-enrichment following gluten exposure when using nucleotide 4-mer feature vectors, and 721 significantly enriched and 690 de-enriched CDR3β sequences when using amino-acid 3-mer feature vectors for CDR3 clustering and sub-repertoire detection. For the CD Gut dataset, the method identified 628 enriched and 619 de-enriched CDR3s when using nucleotide 4-mer feature vectors and 646 enriched 617 de-enriched CDR3s when using amino-acid 3-mer feature vectors. Figure 4a and 4b show the top 20 enriched CDR3βs detected from both datasets when using nucleotide 4-mer feature vectors (for results from amino-acid 3-mer feature vectors see Supplementary Figure 1S). The results of the analyses using nucleotide 4-mers showed high amount of overlap with the results obtained using amino-acid 3-mer level subsequences (Supplementary Figure S2), especially considering the high diversity of immune repertoire datasets and the associated difficulty of representative repertoire sampling. Thus, suggesting that comparable results could be obtained by using either nucleotide or amino acid level subsequence-based feature vectors, although the latter is more computationally expensive.

**Figure 4:**
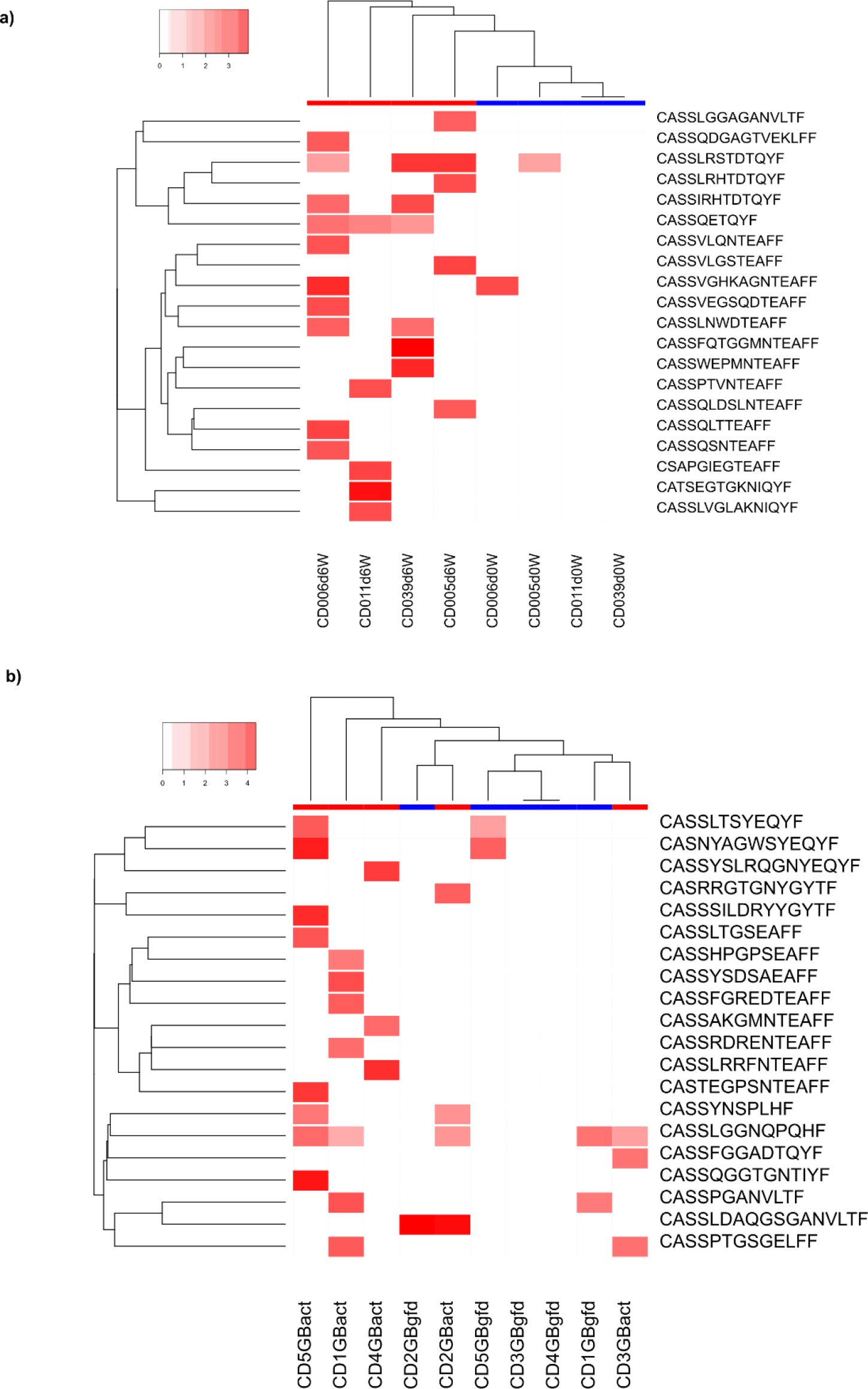
Differentially abundant CDR3β sequences identified by the method using nt 4-mer feature vectors. The top 20 significantly differentially enriched CDR3β sequences during gluten exposure are shown in (a) for CD PBMC and (b) for CD Gut datasets. Low abundance is shown in white and higher abundance is shown in red. The 20 enriched CDR3s are also used to perform hierarchical clustering of the samples with treated samples shown in blue and gluten exposed samples shown in red bars at the top.

To evaluate the overall effectiveness of the method in identifying condition-associated CDR3s, we looked closely at the set of differentially enriched CDR3βs and compared them to known celiac disease associated CDR3β sequences in the literature. The detected lists of enriched CDR3βs had significantly increased usage of previously reported TRBV-genes in gluten-reactive CDR3β sequences (Qiao *et al.*, 2011, 2013; Broughton *et al.*, 2012; Jabri and Sollid, 2017; Han *et al.*, 2013), with the enriched CDR3βs from CD PBMC dataset showing biased usage of TRBV-gene families 4, 7, and 9, specifically TRBV04-02, TRBV07-02 and TRBV09-01 (Figure 5a, and Figure 3Sa), and the enriched CDR3βs from CD gut dataset showing biased usage of TRBV05-08, TRBV06-01, TRBV06-05, TRBV07-05, TRBV07-09, and TRBV10-03 (Figure 5b, and Figure 3Sb). Furthermore, per-position amino acid usage analysis of the enriched CDR3s using some of the over-used TRBV genes provides interesting insights. Besides detecting CDR3βs with the already known amino acid motifs in gluten-reactive CDR3βs with a dominant usage of arginine (R) in position 6 (Qiao *et al.*, 2011, 2013) (Figure 5c and d, Figure 3Sc and d), the method identified other previously unreported over used genes such as TRBV03-01 with previously unappreciated per-position amino acid usage patterns in the detected enriched CDR3s (Figure 5c), the detailed analysis of which may provide insights on their targets and overall significance in the pathogenesis of celiac disease. In addition, except in the nucleotide subsequence based analysis of CD Gut, the list of enriched CDR3βs from both CD PBMC and CD Gut contained significantly high number of previously reported CD-associated CDR3β sequences by Qiao *et al.*, Han *et al.* and Petersen *et al.* (Qiao *et al.*, 2011; Han *et al.*, 2013; Petersen *et al.*, 2014) than was expected by chance, as determined by both using a randomization test or a straight forward comparison to the proportion of previously known celiac disease associated CDR3s in the total repertoire of the combined dataset of all samples (see Tables 2 and 3, see supplementary Tables 1S and 2S for the list of previously known celiac disease associated CDR3 identified by the method). Overall, the method performed well in identifying significant numbers of previously known celiac disease associated CDR3βs, as well as features associated with them, such as bias in v-gene and amino acid usage (see Supplementary dataset 1 for a list of all identified differentially enriched celiac disease associated CDR3β sequences).

**Figure 5:**
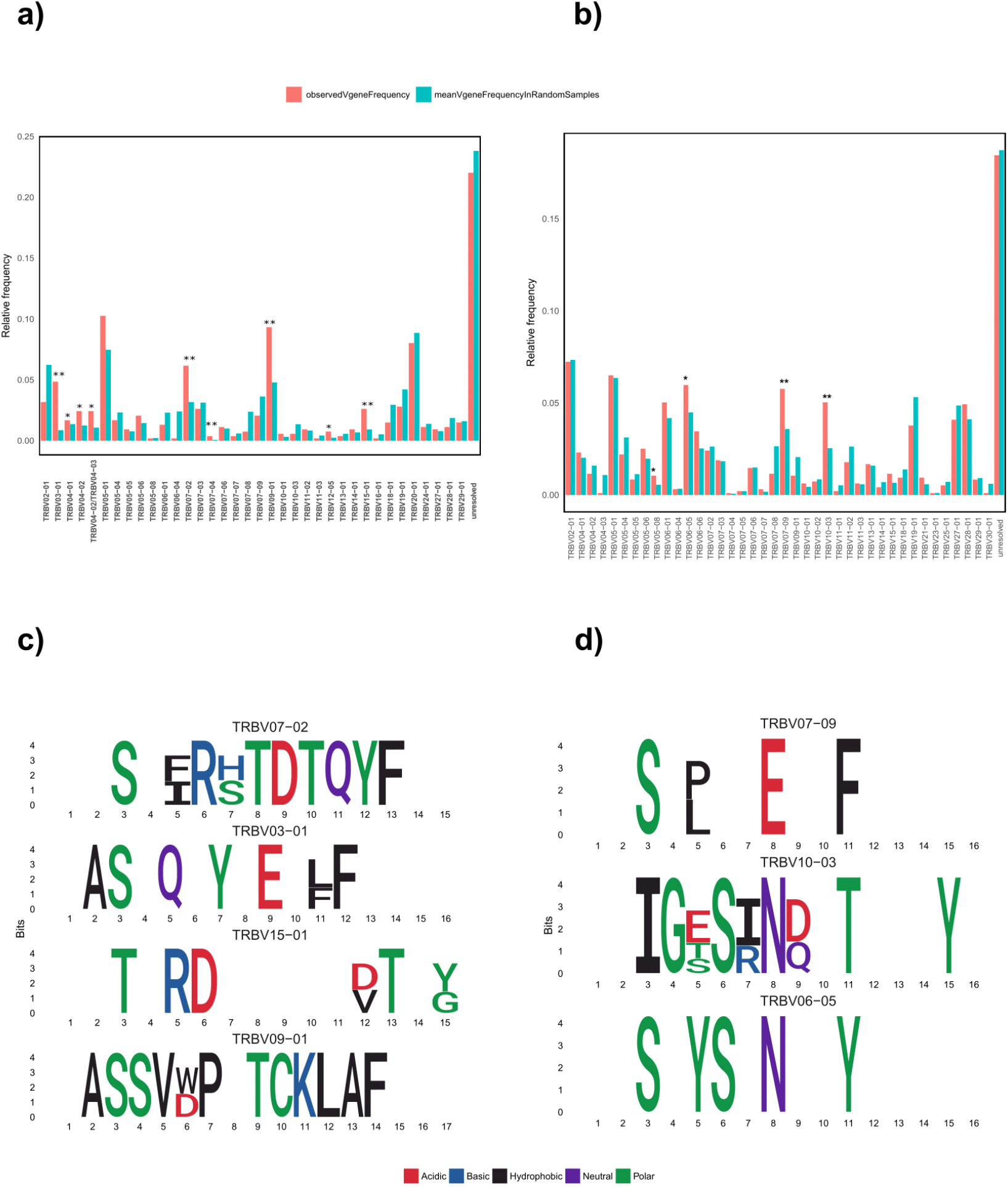
Characteristics of the differentially abundant CDR3β sequences in CD PBMC and CD Gut. The differentially enriched CDR3β sequences had biased usage of TRBV genes that are known to be over-represented in gluten reactive CDR3β sequences in previous studies, such as TRBV07-02 and TRBV09-01 from CD PBMC (a), and TRBV07-09 from CD Gut (b) (observed frequencies are shown in red, mean frequency from randomly generated sets of CDR3s are shown in blue). Significantly, overused amino acids at each position are shown for the enriched CDR3β sequences that use TRBV genes detected to be over-used from CD PBMC (c) and CD Gut (d), amino acids are colored according to their properties. TRBV and per-position amino acid over-usage is assessed by comparing the observed frequencies in the set of differentially enriched CDR3s to that obtained by chance in 100 randomly sampled CDR3s of same size, with p<0.05 considered significant. The results from using nt 4-mer feature vectors are shown.

**Table 2:** Previously known condition-associated CDR3s in the list of DA enriched CDR3s identified by the method (using randomization test)

| Dataset | known condition-associated CDR3s in all samples | Feature space | known condition - associated CDR3s in DA enriched | Permutation test p-value |
| --- | --- | --- | --- | --- |
| CD PBMC | 56 | nt 4-mer | 9 | p=0.0 |
| CD PBMC | 56 | aa 3-mer | 8 | p=0.0 |
| CD GUT | 45 | nt 4-mer | 1 | p=0.6 |
| CD GUT | 45 | aa 3-mer | 4 | p=0.0 |
| YFV PBMC | 7527 | nt 4-mer | 202 | p=0.0 |
| YFV PBMC (1k) | 7527 | aa 3-mer | 80 | p=0.0 |

**Table 3:**
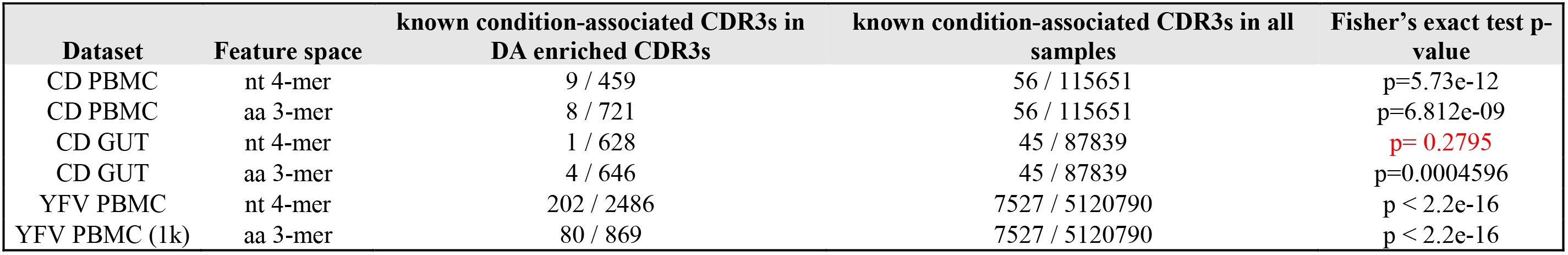
Previously known condition-associated CDR3s in the list of DA enriched CDR3s identified by the method compared to the frequency of known CDR3s in the total combined dataset (using fisher’s exact test)

To assess the performance of the method on publicly available repertoire datasets, we analyzed the yellow fever vaccination CD8+ T-cell CDR3β repertoire dataset (YFV PBMC, n=9, obtained from immuneAccess (DeWitt *et al.*, 2015)) to identify CDR3β sequences responding to the YF-17D vaccine. We compared the total pre-vaccination (day 0) PBMC repertoires to the total post-vaccination (day 14) PBMC repertoires of same individuals. The list of enriched CDR3βs the method identified contained significantly high numbers of vaccine induced CDR3βs from the reported induced day 14 activated effector CD8+ T-Cell CDR3βs in the original publication across all individuals (Table 2 and 3), suggesting that the method can find utility in various study types where detection of T-cell clonotypes with significant expansion is required.

We benchmarked the performance of the method in comparison to four other recently published methods for identification of condition-associated CDR3s. The methods are different in their design and application: our previous published method, Yohannes method (Yohannes *et al.*, 2017) and vdjRec (Pogorelyy, Minervina, Chudakov, *et al.*, 2018) allow comparison at the population level but work only on public CDR3s (and thus identify only public condition associated CDR3s). Conversely, DeWitt’s method (DeWitt *et al.*, 2015) and Alice (Pogorelyy, Minervina, Shugay, *et al.*, 2018) allow comparison only at individual level but allow detection of both private and public CDR3s. Only our current method, also referred to as RepAn, allows population level comparison for detection of both private and public CDR3s. To benchmark the current method, we evaluated how well the methods detect the 56 previously known CD-associated CDR3s that exist in the CD PBMC dataset. Our method (using nt 4-mer kmers) identified 9 of the 56 known CD associated CDR3s as differentially enriched, with these 9 making up 2% of the total 459 CDR3s it identified as CD associated. Alice also detected 9 of the 56, making up 6% of the total of 151 CDR3s it detected as CD-associated. The DeWitt method identified 12 of the 56 known clones among the total of 2003 CDR3s it called as CD associated (0.6%). Our previous method, Yohannes Method, and vdjRec identified 4 and 1 of the 56 known CDR3s, among their 33 (12%) and 31 (3%) CD associated public CDR3s they detected respectively. For the subject level only methods, we collected the union of all identified CDR3s across all individuals as CD associated, this entails that some CDR3s that have contradictory enrichment tendencies in different individuals would be considered as condition associated while population level approaches may not consider such CDR3s as condition-associated. On the other hand, population level methods require a strong signal in multiple subjects to detect condition associated CDR3s, thus may miss CDR3s with low level signal or that are highly private. Given these considerations, our current method performed comparably well, performing much better than the public only methods in the detection of both public and private CD-associated CDR3s, and performing as well as the subject level methods of Dewitt et al., and Alice (Table 4) while allowing population level analysis. The list of the 56 CD-associated CDR3s in the CDPBMC dataset along with the performance of the methods for their detection is listed in supplementary dataset 1.

**Table 4:**
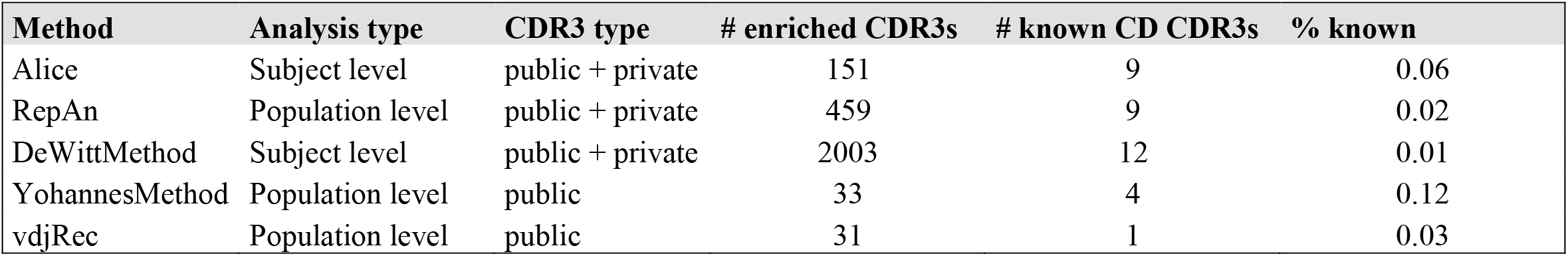
Comparison to other published methods. The current method, RepAn, identifies 9 of the 56 CD-associated CDR3s that exist in the CD PBMC dataset.

| Method | Analysis type | CDR3 type | # enriched CDR3s | # known CD CDR3s | % known |
| --- | --- | --- | --- | --- | --- |
| Alice | Subject level | public + private | 151 | 9 | 0.06 |
| RepAn | Population level | public + private | 459 | 9 | 0.02 |
| DeWittMethod | Subject level | public + private | 2003 | 12 | 0.01 |
| YohannesMethod | Population level | public | 33 | 4 | 0.12 |
| vdjRec | Population level | public | 31 | 1 | 0.03 |

## Discussion

High-throughput immune repertoire sequencing allows in-depth investigation of the adaptive immune profile in healthy and diseased conditions. However, the immense diversity inherent in immune repertoires and inadequate sampling of such diversity, and the small amount of observed repertoire overlap between unrelated individuals makes systematic population level comparison of the adaptive immune response challenging. Thus far, the analyses of disease associated changes in Repseq datasets have mostly been limited to the analysis of repertoire-wide descriptive metrics such as repertoire diversity, repertoire similarity, V-and J-gene segment usage computed at the level of the global repertoires, or the identification condition-associated clonotypes by mostly comparing only the shared or “public” clonotypes across sample groups (Pogorelyy, Minervina, Chudakov, *et al.*, 2018). More specifically, the identification of condition-associated CDR3 sequences is critical in the characterization of the repertoire specific to an antigen and is potentially more valuable in studying disease immunopathology and the overall monitoring of immune repertoires, especially for immediate clinical applications. Thus, as the private response dominates adaptive immune responses, the ability to examine the private response in addition to the public CDR3s is crucial. The computational pipeline we presented in this work allows comparison at the population level to identify both public and private clonotypes associated with conditions, increasing the capacity to detect condition-associated T-cell receptor CDR3 sequences, and thus providing improved ability to monitor changes in the immune repertoire.

We made two main assumptions in our proposed method. Firstly, we assumed that the immune repertoire specific to an antigen would contain T-cell clones with high similarity in their T-cell receptors (TCRs) forming a cluster (or group of clusters) that is distinct, in-terms of TCR sequence, from other T-cells that are not specific to the antigen. This assumption originally stemmed from observations of the celiac disease associated CDR3β sequences in our and other previous studies (Yohannes *et al.*, 2017; Qiao *et al.*, 2011; Dahal-Koirala *et al.*, 2016). Recent works by Dash et.al. and Glanville et.al. (Dash *et al.*, 2017; Glanville *et al.*, 2017) showed that tetramer sorted antigen specific TCRs from different individuals have high sequence similarity and could be grouped into clusters with common specificity to an antigen, further justifying the validity of our assumptions, although just within an antigen specific repertoire. Different ways of representing the CDR3 sequences have been used in recent immune repertoire studies in order to ascertain sequence similarity. We represented CDR3 sequences using a simple, high dimensional subsequence frequency vector, which was then used to define distance in that feature space and cluster CDR3 sequences into similar groups. Greiff *et al.* recently found such immuno-genomic representation of CDR3s to be highly meaningful in allowing the prediction of private versus public CDR3 sequences with high accuracy (Greiff *et al.*, 2017). Although applied for classification of total repertoire samples, Thomas *et al.* used Atchley factors to represent amino-acid subsequences of CDR3 (Thomas *et al.*, 2014; Atchley *et al.*, 2005). On the other hand, direct comparison of the receptor sequences is also possible without CDR3 representation by numeric vectors. Dash *et al.* defined a metric called TCRdist, that uses the amino acid receptor sequences directly, with a weighted Hamming distance of the amino acid sequences of not only the CDR3, but also the CDR1 and CDR2 of both alpha and beta chains to determine the distance between two T-cell receptors (Dash *et al.*, 2017) while Glanville *et al.* combined hamming distance between amino acid CDR3 sequences, with usage patterns of k-mer subsequences in structurally determined positions of high antigen contact propensity to measure distances between pairs of CDR3s (Glanville *et al.*, 2017).

Secondly, we assumed that the clusters of TCRs specific to an antigen encode an important immuno-genomic information in the immune response that is shared across unrelated individuals and could probably be detected from the global repertoire and across treatment conditions. Using a feature space of 4-mer nucleotide or 3-mer amino acid subsequence frequencies to represent each CDR3 sequence and dissecting total immune repertoires into units or sub-repertoires that exist across individuals, our method successfully identified previously reported and new condition-associated CDR3β sequences (both private and public) from the datasets we analyzed, demonstrating the validity of our assumption.

Evaluating the effectiveness of the method developed in this study, especially its sensitivity and specificity, requires baseline datasets, conceivably simulated, with an exhaustive list of condition-associated CDR3s. Real or simulated datasets of the adaptive immune repertoire are unavailable, and simulating an adaptive response scenario is highly complicated due to our limited understanding of TCRs involved in immune responses. Thus, we simply evaluated the method’s effectiveness by comparing the features of the list of differentially enriched condition-associated CDR3s obtained to known antigen binding condition-associated CDR3s from previous studies. Our results showed that the list of putative condition-associated CDR3s the method produced contained statistically significant numbers of known condition-associated CDR3s (both celiac disease and yellow fever vaccination datasets) than could be obtained by chance and have significant enrichment of V-gene and per-position amino acid usage typical of known condition-associated CDR3s, validating the high usability of the proposed computational pipeline. We also benchmarked the performance of the method compared to recently published similar methods in detecting 56 previously reported gluten-specific CDR3 sequences present in CD PBMC data. The method performed comparably well and identified 9 of the 56 known CDR3s, only next to the method by DeWitt *et. al*, which works at the subject level and requires multiple samples from an individual, and as well as Alice which works at the subject level. Although Alice performed better than all methods in terms of percentage of known CDR3s among total detected private and public CDR3s, the total number of condition-associated CDR3s it detected was smaller than both the current method, and DeWitt *et.al*’s method. Since identifying more number of relevant CDR3s is also an important goal of such methods, and the current method performed second only to DeWitt et.al’s method in that regard, the method provides an intermediate performance between the highly conservative Alice and the highly exhaustive DeWitt *et.al*’s method, while also performing as good in identifying the known gluten-specific CDR3s. In addition, as the only population level method that allows detection of both private and public CDR3s, it has the added advantage of being usable in the direct comparison of groups of immune repertoires, and thus the only current method suited for study designs that intend to compare repertoires at the population level between samples in different conditions.

Condition-associated TCR CDR3s are not fully known for many multifactorial autoimmune or cancer related diseases. For diseases with known antigen, such as celiac disease (CD) where some information about the antigen (gluten) is known, specific protocols like tetramers and/or sorting would have to be designed to characterize the antigen-specific T-cells, their CDR3s, and other phenotypes in detail. Although highly useful, such methods designed to select antigen-specific repertoires are unable to detect T-cell clones responding to other important immune targets other than gluten, either self or foreign, possibly ignoring a crucial part of the immune response that would explain the pathology even more. Methods such as the one presented in this study that attempt to detect T-cell clones from total repertoires that show significant association with a condition, without necessarily having prior knowledge of the immune target (or targets), coupled with techniques that profile the overall gene expression of the condition-associated T-cell clones, would potentially provide a highly comprehensive picture of the adaptive immune response. This leads to a much more complete understanding of such diseases in-terms of unraveling the hidden pieces of the puzzle, such as by helping identify the unknown immune targets. For diseases for which the antigens are totally unknown, the application of such methods could lead to the identification of the associated CDR3 sequences that could serve as immune-response bio-markers with possible clinical application, as well as enabling their comparison to other known disease associated CDR3 sequences available in CDR3 databases (Tickotsky *et al.*, 2017; Shugay *et al.*, 2018), potentially allowing prediction of their target or antigen types.

Main limitations of the methodology include restricted repertoire representativeness and computational intensiveness, both arising from the immense diversity of immune repertoires. There are millions of unique CDR3 sequences in every person, each representing a T-cell clone, and most found only in a single person. Only tens to hundreds of thousands of unique CDR3s are being sampled with the current Repseq technology per sample. Since calculating pairwise distances for potentially hundreds of thousands of CDR3s is computationally intensive, if not infeasible, we adopted repeat resampling and applied the methodology a repeated number times with randomly selected smaller repertoire samples. The number of repeat resample runs chosen and the repertoire size of each repertoire resamples determines the computational resources required and comprehensiveness of the result.

To conclude, by clustering CDR3 sequences into groups with similar immunogenomic features, and finding their close matches across different samples, we showed that condition-associated CDR3 sequences that are private or public, and with significant differential abundance, can be detected at the population level. The approach paves the way for the identification of private or public CDR3s (and their features) associated with diseases or other important phenotypes such as HLA-type, possibly allowing comprehensive categorization and archiving of T-cell clonotypes, which has a vast potential in understanding the adaptive immune response in various disease conditions and disease development stages, identifying unknown self or foreign antigens in diseases with unknown immune targets, examining immunological history encoded in the immune repertoire, and possible early prediction of the adaptive immune response.

## Supporting information

## Acknowledgments

We thank Anne Heimonen, Marja-Terttu Oksanen, Hanne Ahola and Andrea de Kauwe for their help in patient recruitment, sample collection and handling, and laboratory work. We also thank Rigbe G. Weldatsadik for helpful discussions and proof reading of the manuscript. We acknowledge CSC – IT Center for Science, Finland, for providing computational resources.

## Author contributions

Study concept and design: DAY, DG, PS; acquisition of study samples, technical and material support: KK, KK, PS; acquisition of data: PS, DAY; analysis and interpretation of data: DAY, PS, DG; computational pipeline design and statistical analysis: DAY, DG; pipeline implementation: DAY, DG; manuscript drafting: DAY, DG, PS; critical revision of manuscript: DG, PS; Study supervision: PS & DG.

## Supplementary Materials

**Figure 1S:**
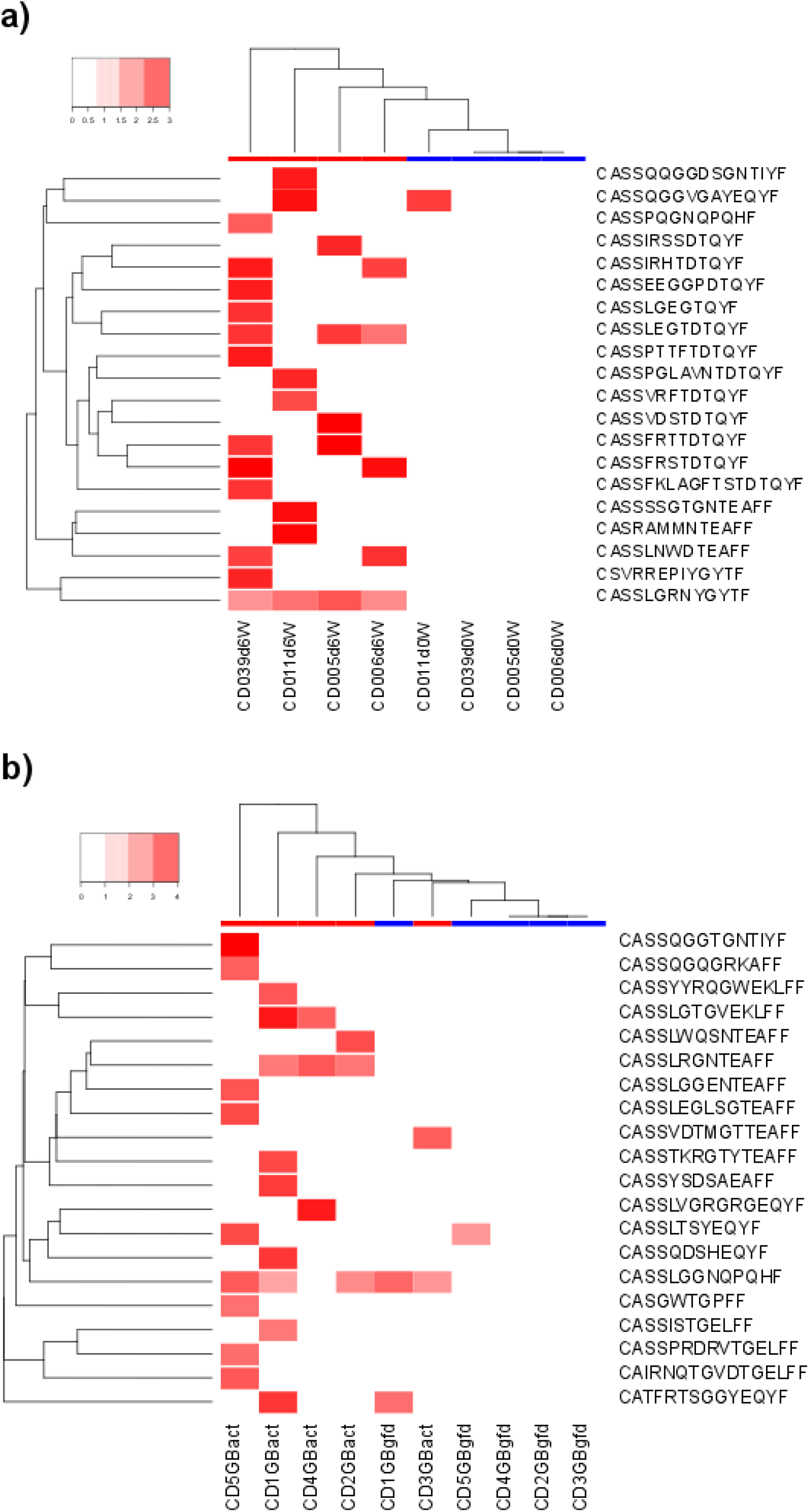
Differentially abundant CDR3b sequences in CD PBMC and CD Gut. (a) and (b) show top 20 significantly differentially enriched CDR3b sequences during gluten exposure in CD PBMC and CD Gut datasets respectively from the DA analysis using aa 3-mer feature vectors.

**Figure 2S:**
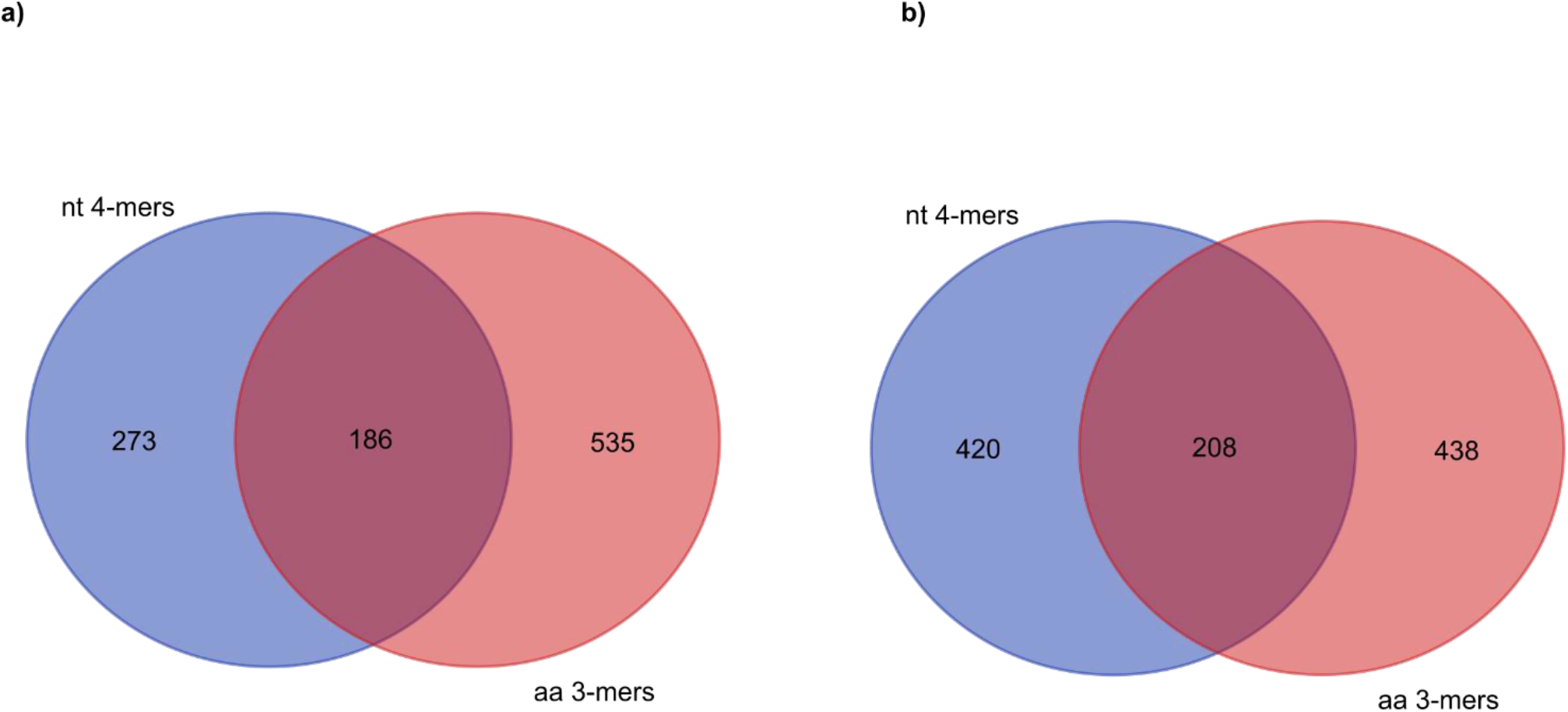
The overlap between the differentially enriched CDR3β sequences of the DA analyses using nt 4-mer and aa 3-mer feature vectors is shown for (a) CD PBMC and (b) CD Gut datasets.

**Figure 3S:**
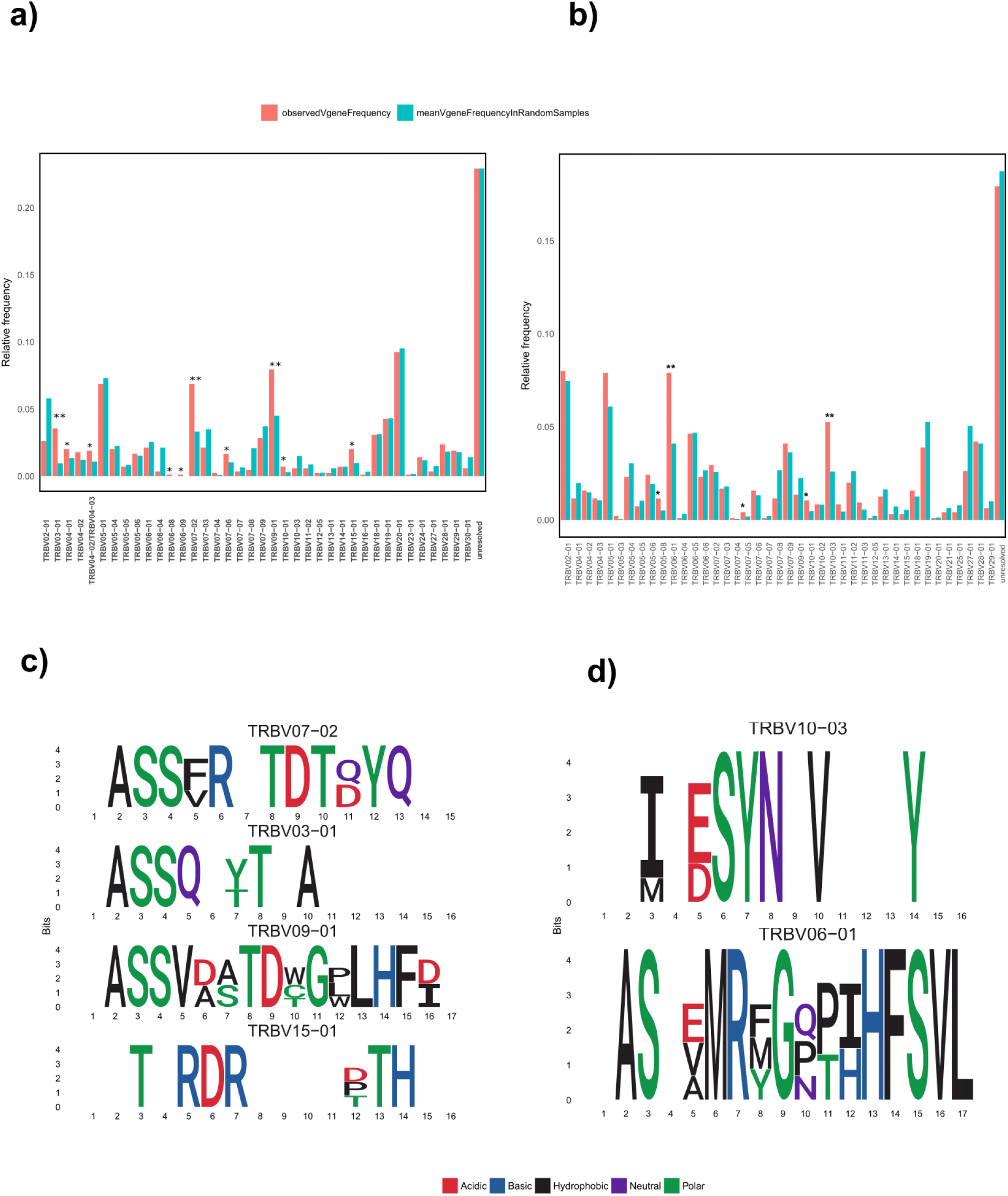
Characteristics of the differentially abundant CDR3b sequences in CD PBMC and CD Gut. The differentially enriched CDR3b sequences had biased usage of TRBV genes that are known to be over-represented in gluten reactive CDR3b sequences in previous studies, such as TRBV07-02 and TRBV09-01 from CD PBMC (a), and TRBV07-09 from CD Gut (b) (observed frequencies are shown in red, mean frequency from randomly generated sets of CDR3s are shown in blue). Significantly over-used amino acids at each position are shown for the enriched CDR3b sequences that use TRBV genes detected to be over-used from CD PBMC (c) and CD Gut (d), amino acids are colored according to their properties. TRBV and per-position amino acid over-usage is assessed by comparing the observed frequencies in the set of CDR3s to that obtained by chance in 100 randomly sampled CDR3s of same size, with p<0.05 considered significant. The results from using aa 3-mer feature vectors are shown.

**Table 1S:**
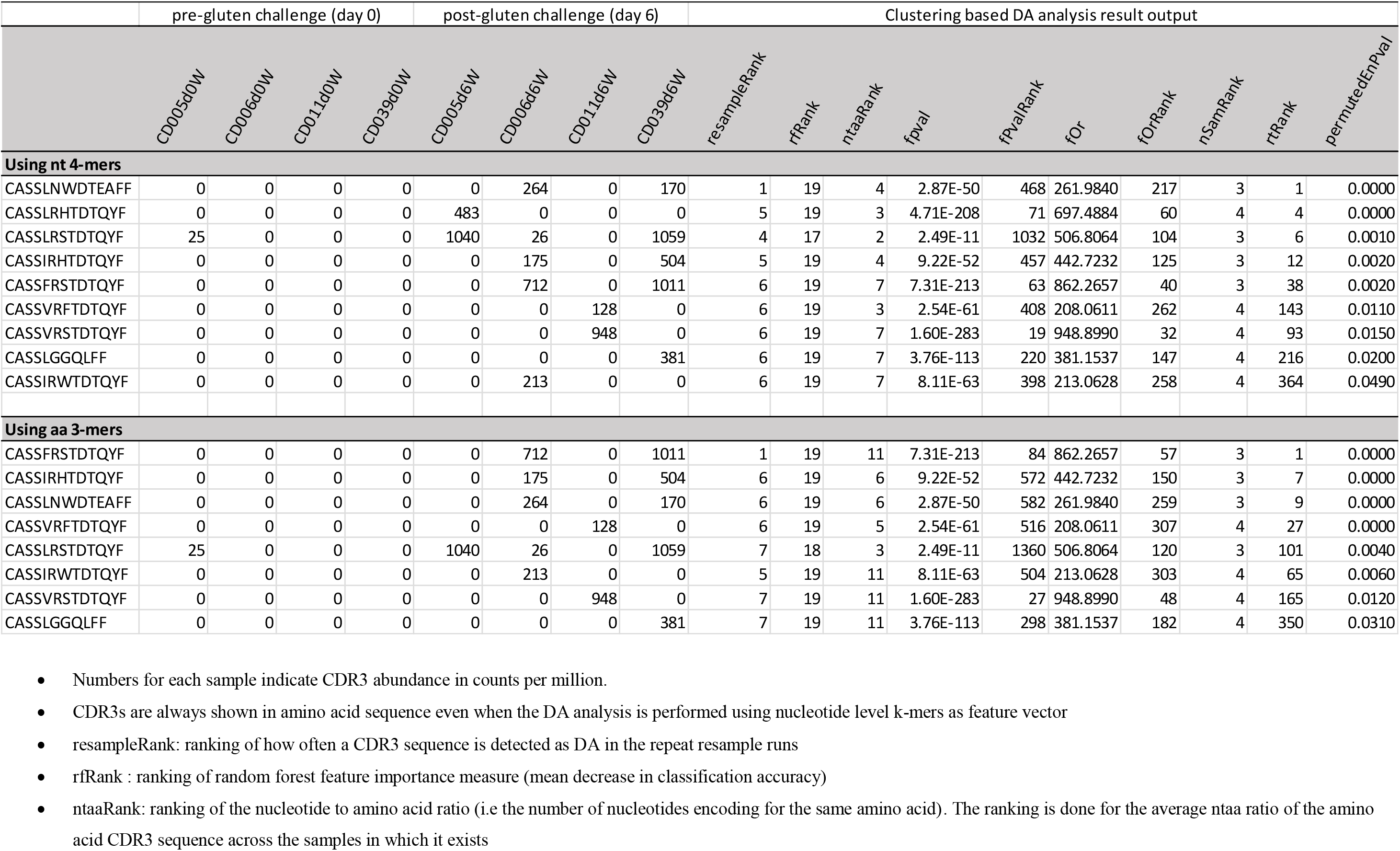
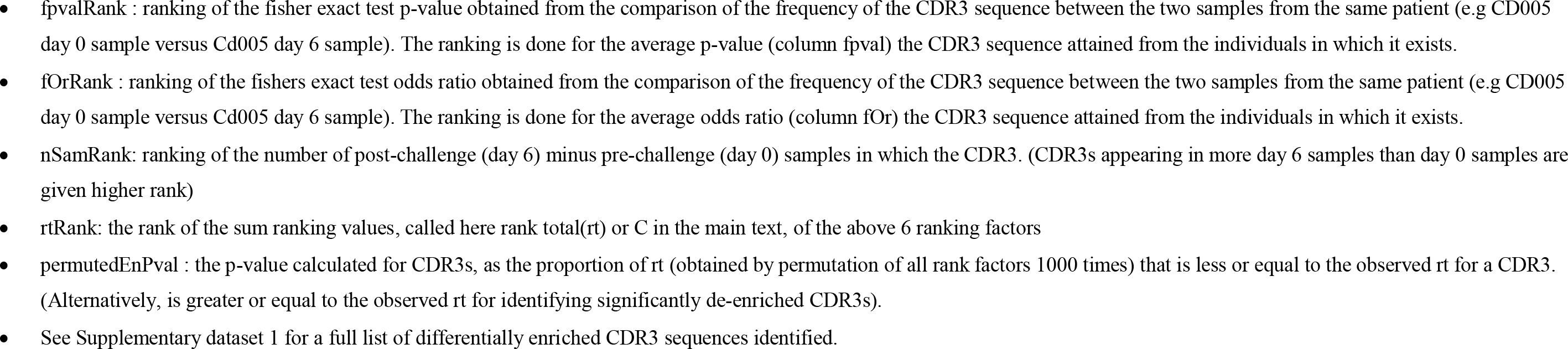
List of previously known gluten-reactive celiac disease associated CDR3b sequences identified as differentially enriched by the method from the CD PBMC repertoire dataset.

**Table 2S:**
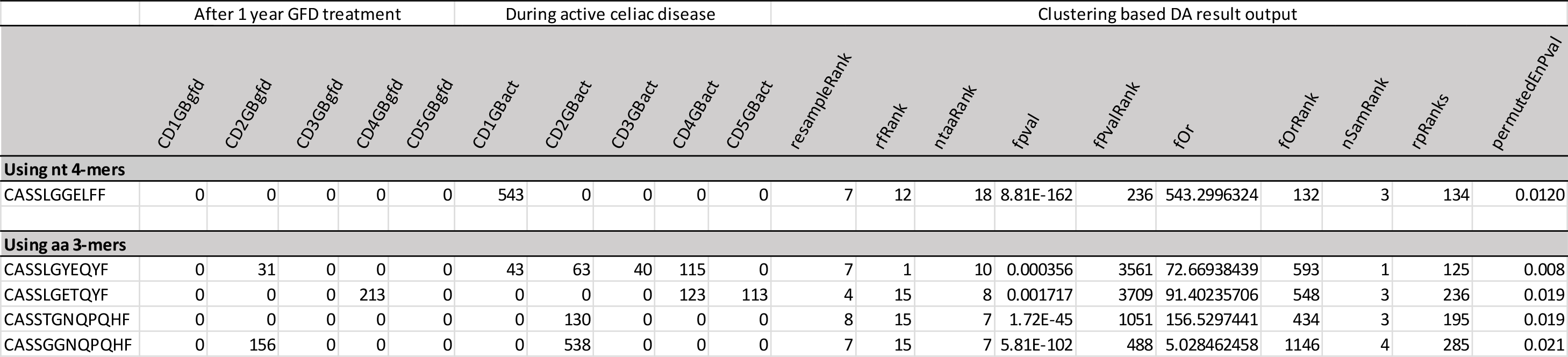
List of previously known gluten-reactive celiac disease associated CDR3b sequences identified as differentially enriched by the method from the CD Gut repertoire dataset.

**Figure 4S:**
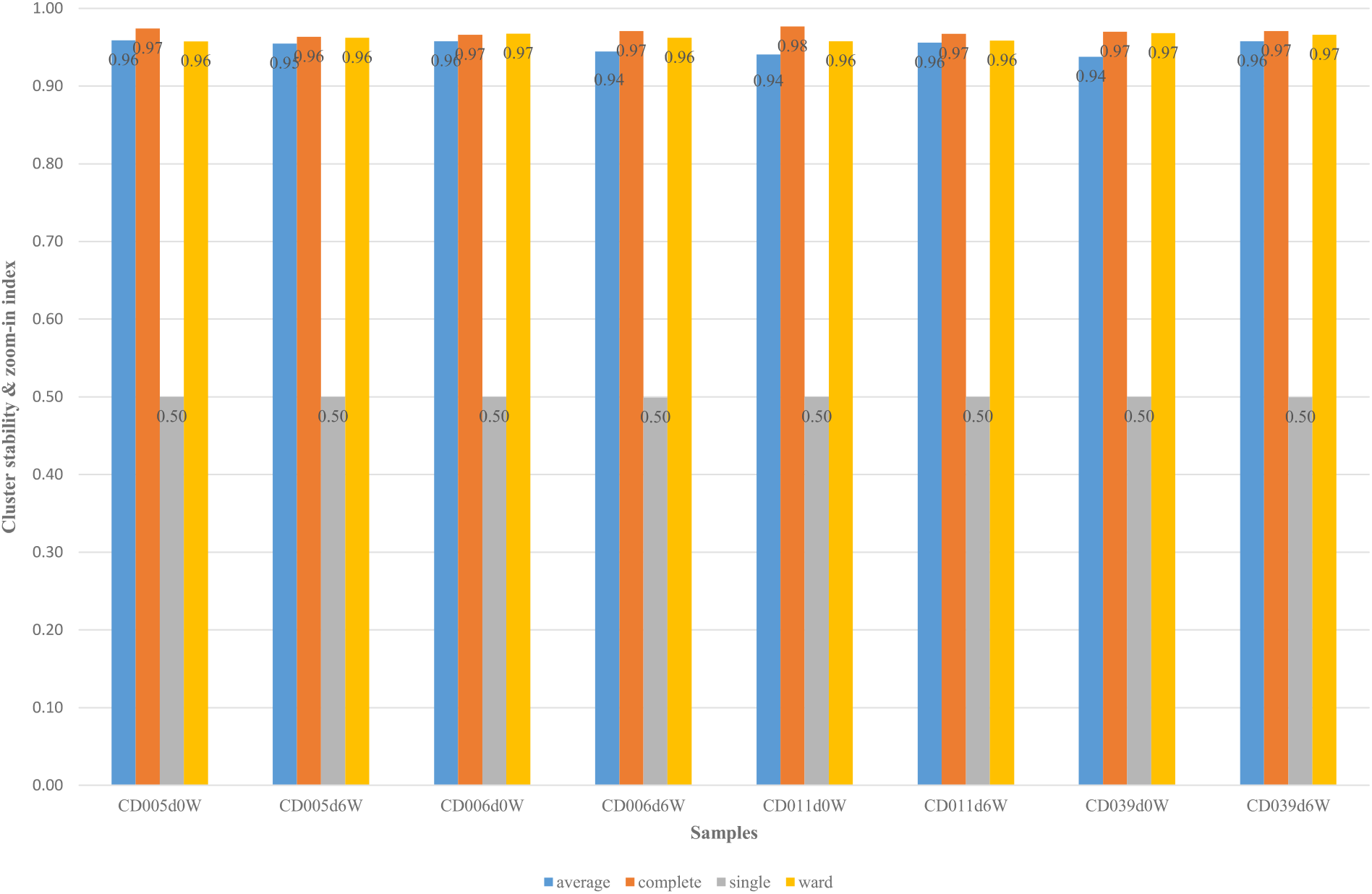
Comparison of linkage methods in the hierarchical clustering of TCR CDR3s. Linkage methods were evaluated using a combined measure of cluster stability (the cls.stab.sim.ind in R package clv (Nieweglowski, 2013), using the Rand similarity index), and a zoom-in factor (1-(average cluster size)/(total number of CDR3s)), favoring smaller sized clusters relative to the total number of starting CDR3s that allow deeper zooming into the diverse repertoire samples. We sum the two values and divide by 2 to get a stability & zoom-in index between 0 and 1, with 1 meaning stable clustering with high number of small sized clusters. We resampled 5000 unique TCR CDR3 sequences from all CD PBMC samples and evaluated the clustering performance of commonly used linkage methods in hierarchical clustering: average, complete, ward, and included the single linkage method. In each case, we performed hierarchical clustering of the CDR3s, partitioned the CDR3s into k clusters as determined by the dynamic tree cut algorithm (Langfelder *et al.*, 2008), and evaluated stability at the determined k number of clusters. The complete linkage method performed consistently better for all samples followed by the ward method. We chose to use the complete method because it gave good stability with more number of clusters, allowing for deeper “zooming-in”, which is critical as it reduces repertoire diversity more.

## Reference

Agathangelidis, A. et al. (2012) Stereotyped B-cell receptors in one-third of chronic lymphocytic leukemia: a molecular classification with implications for targeted therapies. Blood, 119, 4467-4475.

Atchley, W.R. et al. (2005) Solving the protein sequence metric problem. Proc. Natl. Acad. Sci. U. S. A., 102, 6395-6400.

Benati, D. et al. (2016) Public T cell receptors confer high-avidity CD4 responses to HIV controllers. J. Clin. Invest., 126, 2093-2108.

Benichou, J. et al. (2012) Rep-Seq: uncovering the immunological repertoire through next-generation sequencing. Immunology, 135, 183-191.

Breitling, R. et al. (2004) Rank products: a simple, yet powerful, new method to detect differentially regulated genes in replicated microarray experiments. FEBS Lett., 573, 83-92.

Broughton, S.E. et al. (2012) Biased T cell receptor usage directed against human leukocyte antigen DQ8-restricted gliadin peptides is associated with celiac disease. Immunity, 37, 611-621.

Covacu, R. et al. (2016) System-wide Analysis of the T Cell Response. Cell Rep., 14, 2733-2744.

Dahal-Koirala, S. et al. (2016) TCR sequencing of single cells reactive to DQ2.5-glia-α2 and DQ2.5-glia-ω2 reveals clonal expansion and epitope-specific V-gene usage. Mucosal Immunol., 9, 587-596.

Darzentas, N. and Stamatopoulos, K. (2013) Stereotyped B cell receptors in B cell leukemias and lymphomas. Methods Mol. Biol. Clifton NJ, 971, 135-148.

Dash, P. et al. (2017) Quantifiable predictive features define epitope-specific T cell receptor repertoires. Nature.

DeWitt, W.S. et al. (2015) Dynamics of the cytotoxic T cell response to a model of acute viral infection. J. Virol., 89, 4517-4526.

Emerson, R.O. et al. (2017) Immunosequencing identifies signatures of cytomegalovirus exposure history and HLA-mediated effects on the T cell repertoire. Nat. Genet., 49, 659-665.

Glanville, J. et al. (2017) Identifying specificity groups in the T cell receptor repertoire. Nature, 547, 94-98.

Greiff, V. et al. (2017) Learning the High-Dimensional Immunogenomic Features That Predict Public and Private Antibody Repertoires. J. Immunol., 199, 2985-2997.

Han, A. et al. (2013) Dietary gluten triggers concomitant activation of CD4+ and CD8+ αβ T cells and 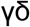 T cells in celiac disease. Proc. Natl. Acad. Sci. U. S. A., 110, 13073-13078.

Ho, T.K. (1995) Random decision forests. In, Document analysis and recognition, 1995. proceedings of the third international conference on. IEEE, pp. 278-282.

Jabri, B. and Sollid, L.M. (2017) T Cells in Celiac Disease. J. Immunol. Baltim. Md 1950, 198, 3005-3014.

Langfelder, P. et al. (2008) Defining clusters from a hierarchical cluster tree: the Dynamic Tree Cut package for R. Bioinforma. Oxf. Engl., 24, 719-720.

Lefranc, M.-P. et al. (2015) IMGT®, the international ImMunoGeneTics information system^®^ 25 years on. Nucleic Acids Res., 43, D413-422.

Li, H. et al. (2012) Determinants of public T cell responses. Cell Res., 22, 33-42.

Liaw, A. and Wiener, M. (2002) Classification and regression by randomForest. R News, 2, 18-22.

Madi, A. et al. (2014) T-cell receptor repertoires share a restricted set of public and abundant CDR3 sequences that are associated with self-related immunity. Genome Res., 24, 1603-1612.

Petersen, J. et al. (2014) T-cell receptor recognition of HLA-DQ2-gliadin complexes associated with celiac disease. Nat. Struct. Mol. Biol., 21, 480–488.

Pogorelyy, M.V., Minervina, A.A., Shugay, M., et al. (2018) Detecting T-cell receptors involved in immune responses from single repertoire snapshots. bioRxiv, 375162.

Pogorelyy, M.V., Minervina, A.A., Chudakov, D.M., et al. (2018) Method for identification of condition-associated public antigen receptor sequences. eLife, 7, e33050.

Qi, Q. et al. (2014) Diversity and clonal selection in the human T-cell repertoire. Proc. Natl. Acad. Sci., 111, 13139-13144.

Qiao, S.-W. et al. (2013) Biased usage and preferred pairing of α- and β-chains of TCRs specific for an immunodominant gluten epitope in coeliac disease. Int. Immunol.

Qiao, S.-W. et al. (2011) Posttranslational modification of gluten shapes TCR usage in celiac disease. J. Immunol., 187, 3064-3071.

Risnes, L.F. et al. (2018) Disease-driving CD4+ T cell clonotypes persist for decades in celiac disease. J. Clin. Invest., 128, 2642-2650.

Robins, H.S. et al. (2009) Comprehensive assessment of T-cell receptor beta-chain diversity in alphabeta T cells. Blood, 114, 4099-4107.

Shugay, M. et al. (2018) VDJdb: a curated database of T-cell receptor sequences with known antigen specificity. Nucleic Acids Res., 46, D419-D427.

Thomas, N. et al. (2014) Tracking global changes induced in the CD4 T-cell receptor repertoire by immunization with a complex antigen using short stretches of CDR3 protein sequence. Bioinforma. Oxf. Engl., 30, 3181-3188.

Tickotsky, N. et al. (2017) McPAS-TCR: a manually curated catalogue of pathology-associated T cell receptor sequences. Bioinforma. Oxf. Engl., 33, 2924-2929.

Vanhanen, R. et al. (2016) T cell receptor diversity in the human thymus. Mol. Immunol., 76, 116-122.

Venturi, V. et al. (2008) The molecular basis for public T-cell responses? Nat. Rev. Immunol., 8, 231-238.

Yohannes, D.A. et al. (2017) Deep sequencing of blood and gut T-cell receptor β-chains reveals gluten-induced immune signatures in celiac disease. Sci. Rep., 7, 17977.

## Reference

Langfelder, P. et al. (2008) Defining clusters from a hierarchical cluster tree: the Dynamic Tree Cut package for R. Bioinforma. Oxf. Engl., 24, 719–720.

Nieweglowski, L. (2013) clv: Cluster Validation Techniques.

